## Supplementary Figures for "Characterisation of a nucleo-adhesome"

**Supplementary Information**

Supplementary Figure 1
Supplementary Figure 2
Supplementary Figure 3
Supplementary Figure 4
Supplementary Figure 5
Supplementary Table 1
Supplementary Table 2
Supplementary Table 3
Supplementary Table 4
Supplementary Table 5
Supplementary Table 6
Supplementary Table 7
Supplementary Table 8
Supplementary Table 9
Supplementary Table 10

[1]Cancer Research UK Edinburgh Centre, Institute of Genetics and Cancer, University of Edinburgh, Edinburgh EH4 2XR, UK. [2]Leiden University Medical Center, 2333 ZC Leiden, The Netherlands. [3]MRC Human Genetics Unit, Institute of Genetics and Cancer, University of Edinburgh, Edinburgh EH4 2XU, UK. [4]Present address: Instituto de Biomedicina y Biotecnología de Cantabria (IBBTEC), Consejo Superior de Investigaciones Científicas (CSIC)-Universidad de Cantabria, Santander, 39011, Spain. [5]Present address: Almac Diagnostic Services, 19 Seagoe Industrial Estate, Craigavon BT63 5QD, UK. [6]Present address: NanoString Technologies, Inc., Seattle, Washington 98109, USA. *These authors contributed equally to this work. [†]

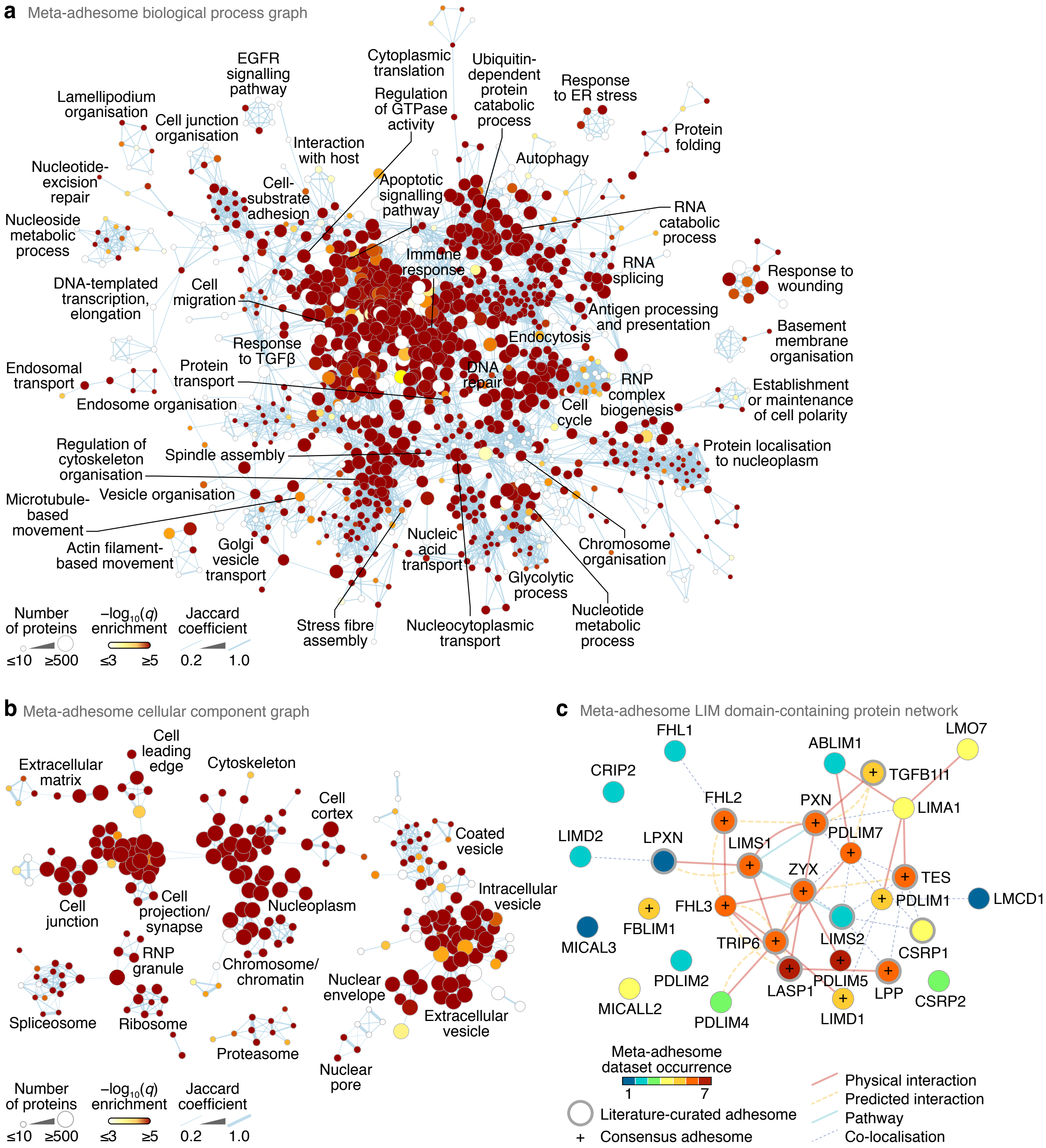


**Supplementary Figure 1.** Functional network analysis of the meta-adhesome. (**a**, **b**) Graph-based clustering of over-represented biological processes (**a**) and cellular components (**b**) in the meta-adhesome. Edge-weighted force-directed graphs were generated from gene-set memberships of enriched Gene Ontology terms ($P < 0.01$, hypergeometric test with Benjamini–Hochberg correction). Connected components with more than five nodes (enriched terms) are shown. Representative functional categories summarising selected clusters of enriched terms are labelled. Node (circle) size represents the number of meta-adhesome proteins annotated for each enriched term; node fill colour represents the significance of enrichment. Edge (line) weight is proportional to the overlap of gene-set membership of connected nodes (Jaccard coefficient ≥ 0.2). Selected clusters are indicated in Fig. 1c, d. (**c**) Functional association network of LIM domain-containing proteins in the meta-adhesome ($P = 6.04 \times 10^{-8}$, hypergeometric test with Benjamini–Hochberg correction). Node fill colour represents frequency of occurrence in meta-adhesome datasets; thick node borders indicate representation in the literature-curated adhesome. Edges represent reported associations. Proteins (nodes) are annotated with gene names for clarity.

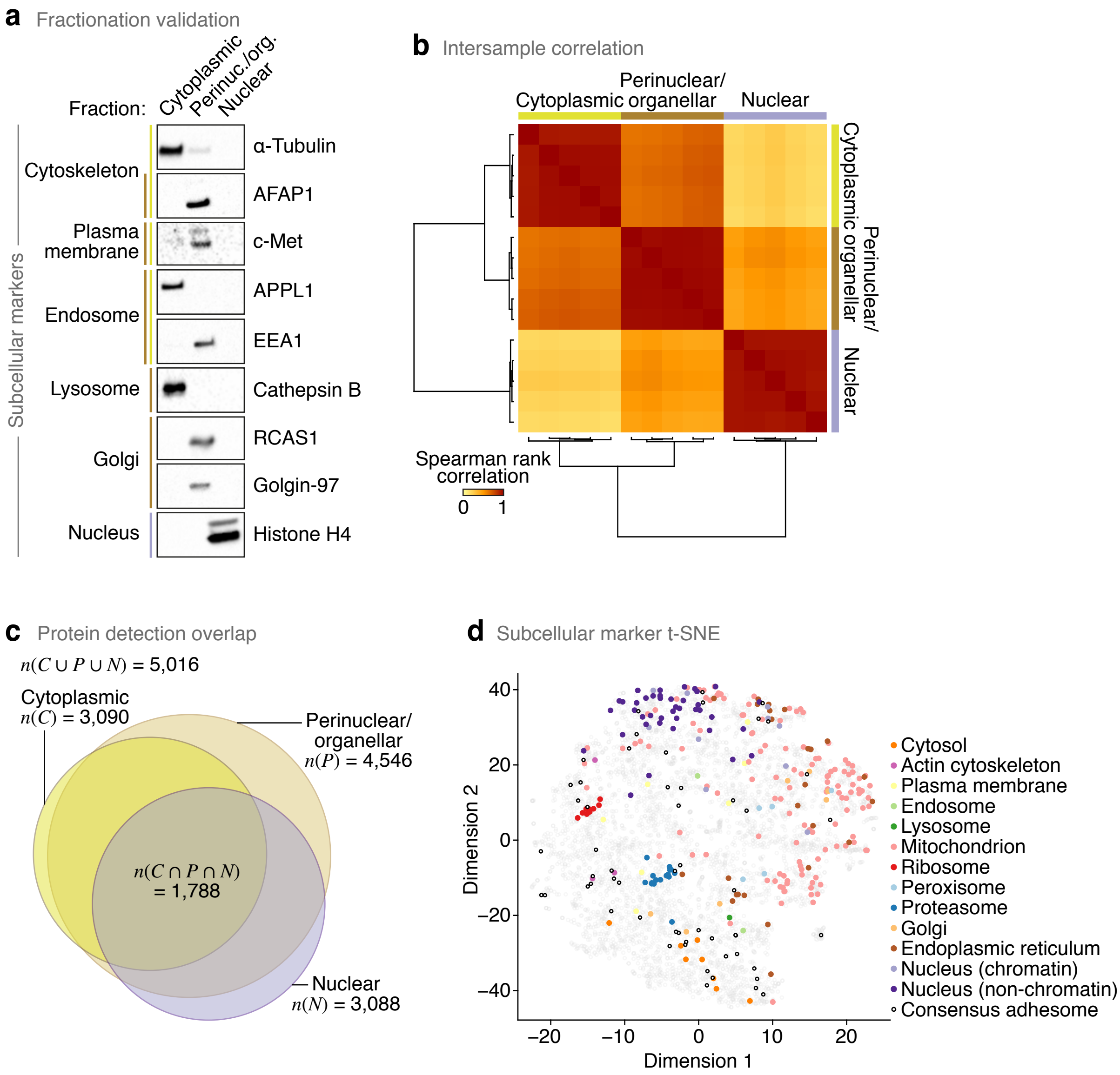


**Supplementary Figure 2.** Mass spectrometric analysis of the subcellular proteomes of SCC cells. (**a**) Subcellular fractionation of SCC cells. Markers for given subcellular locations are indicated. See also Fig. 2b. (**b**) Hierarchical cluster analysis of Spearman rank correlation coefficients for all pairwise sample comparisons based on proteins quantified by LC-MS/MS in at least four out of five biological replicates. (**c**) Area-proportional Euler diagram of numbers of proteins detected and quantified in different subcellular fractions. Cardinalities of selected sets are defined, where $C$ = cytoplasmic, $P$ = perinuclear/organellar and $N$ = nuclear fractions. (**d**) t-SNE map of the subcellular proteomes annotated with curated subcellular markers. Detailed subcellular classification of markers is shown. See also Fig. 2g.

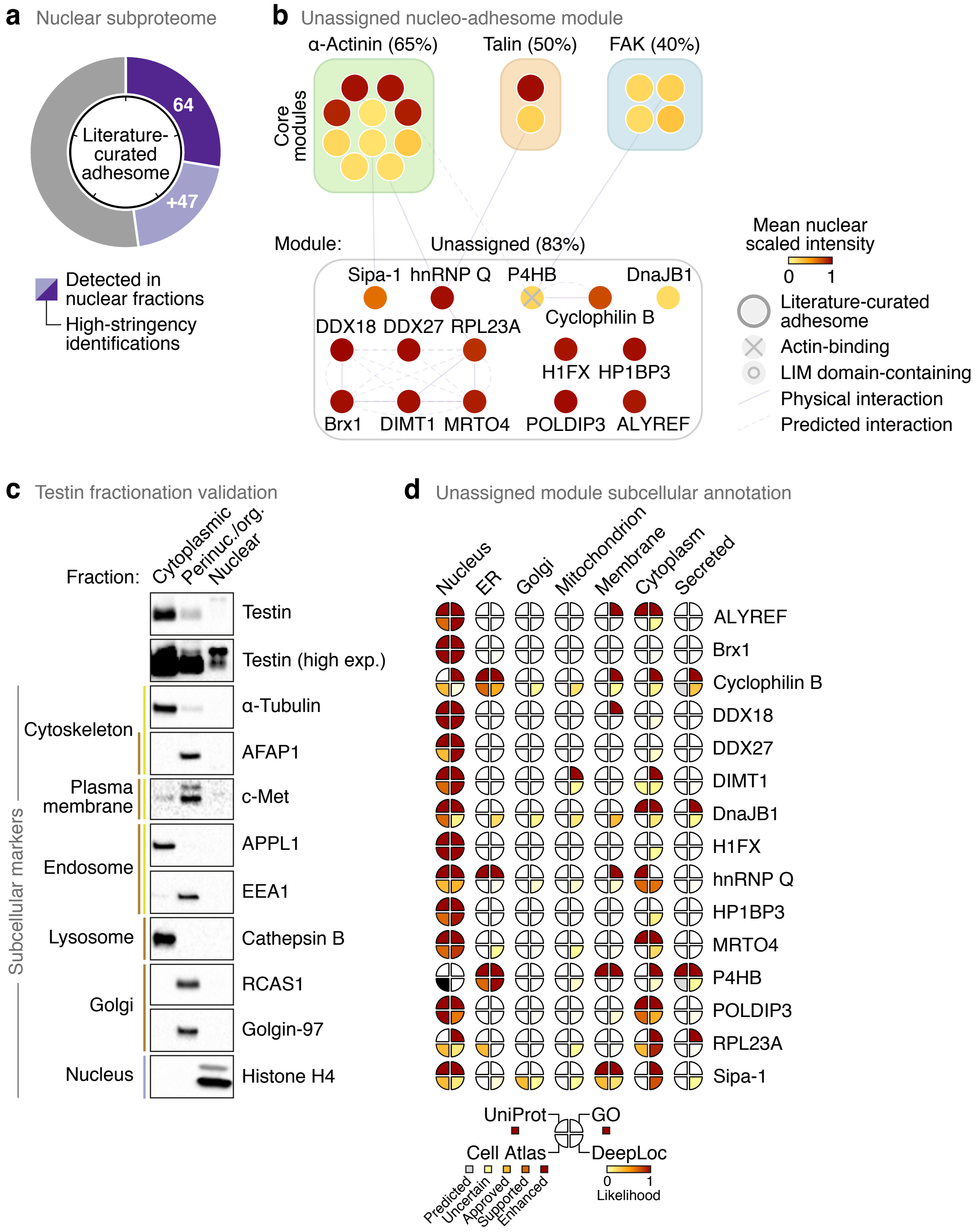


**Supplementary Figure 3.** Network analysis of a nucleo-adhesome. (**a**) Proportion of the literature-curated adhesome quantified in SCC cell nuclear fractions. High-stringency proteins (dark purple segments) were identified in all five biological replicate experiments and quantified with a >5% fraction of the cellular pool; light purple segments indicate additional proteins detected in nuclear fractions. Tick marks indicate 20% increments. (**b**) Network model of high-stringency nuclear proteins present in the consensus adhesome but not associated with an assigned module (termed unassigned). Coverage of the module is indicated in parentheses. Node (circle) fill colour represents the mean scaled intensity of nuclear fraction replicates; thick grey node borders indicate representation in the literature-curated adhesome. Edges (lines) represent reported interactions. (**c**) Subcellular fractionation of SCC cells. Markers for given subcellular locations are indicated. (**d**) Subcellular annotation plot of high-stringency nuclear proteins in the unassigned nucleo-adhesome module. Segments are coloured according to annotation in the UniProt knowledgebase, annotation in the Gene Ontology (GO) cellular component domain, Cell Atlas reliability class and DeepLoc prediction likelihood as indicated. For Cell Atlas annotation, nucleus represents the highest-scoring reliability class of annotated nuclear subregions (e.g. nucleoli, nucleoplasm, nuclear membrane); black segment indicates annotation in mouse cells only (no corresponding reliability class).

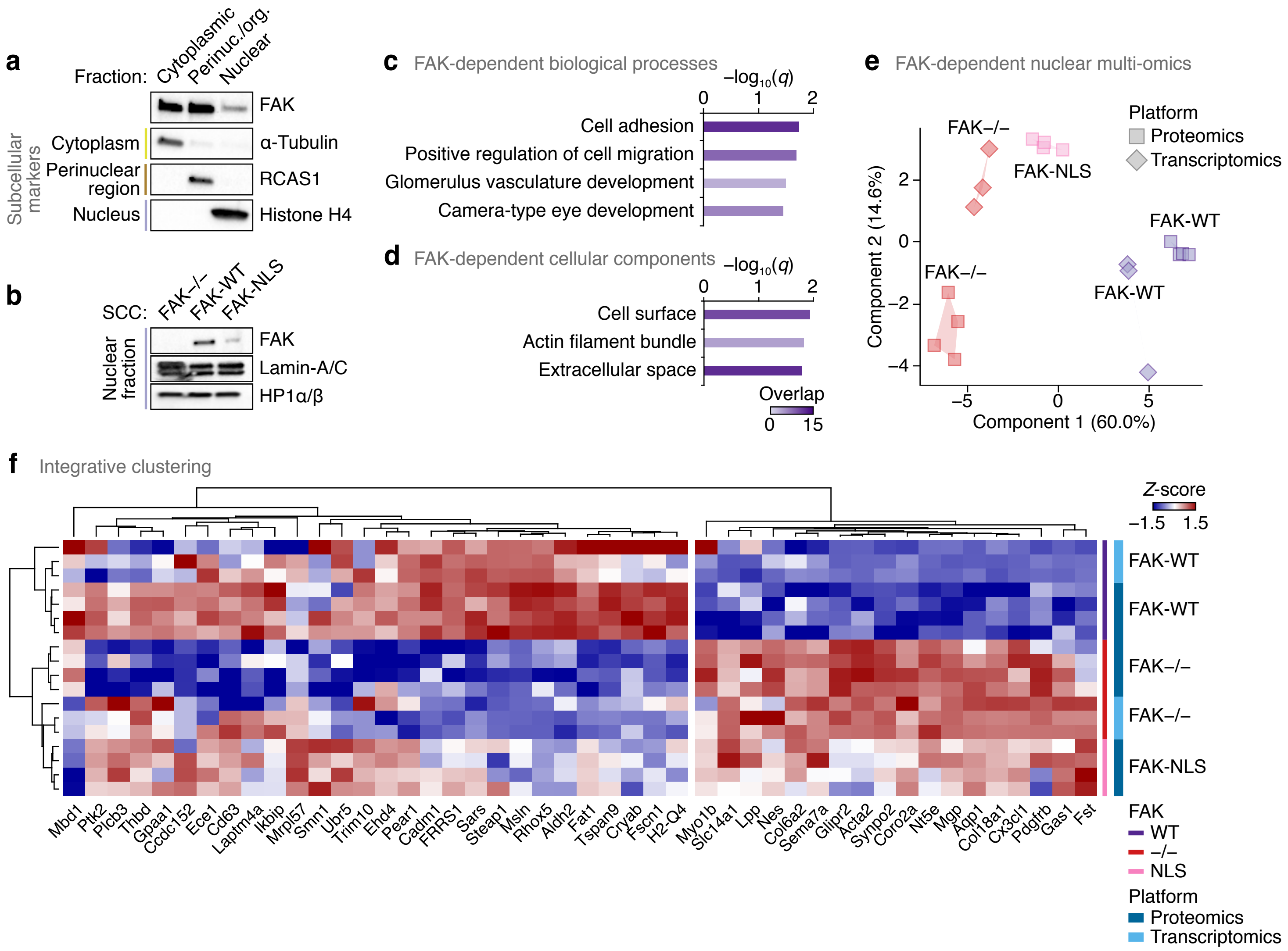


**Supplementary Figure 4.** Mass spectrometric analysis of the FAK-dependent nuclear proteome. (**a**) Subcellular fractionation of SCC cells. Markers for given subcellular locations are indicated. (**b**) Nuclear fractionation of SCC cells that express FAK-WT or FAK-NLS or do not express FAK (FAK–/–). Lamin-A/C and heterochromatin protein 1 α/β (HP1α/β) are nuclear markers. (**c**, **d**) Over-representation analysis of Gene Ontology biological processes (**c**) and cellular components (**d**) in the FAK-dependent nuclear proteome. Purple shading intensity indicates size of subproteome overlap with respective gene sets. Representative terms were determined using affinity propagation. (**e**) Principal component analysis of integrated FAK-dependent nuclear proteomic and FAK-dependent transcriptomic (multi-omic) data. (**f**) Integrative cluster analysis of the FAK-dependent nuclear multi-omic data. Proteins are labelled with gene names for clarity. Sample (row) dendrogram also shown in Fig. 4i.

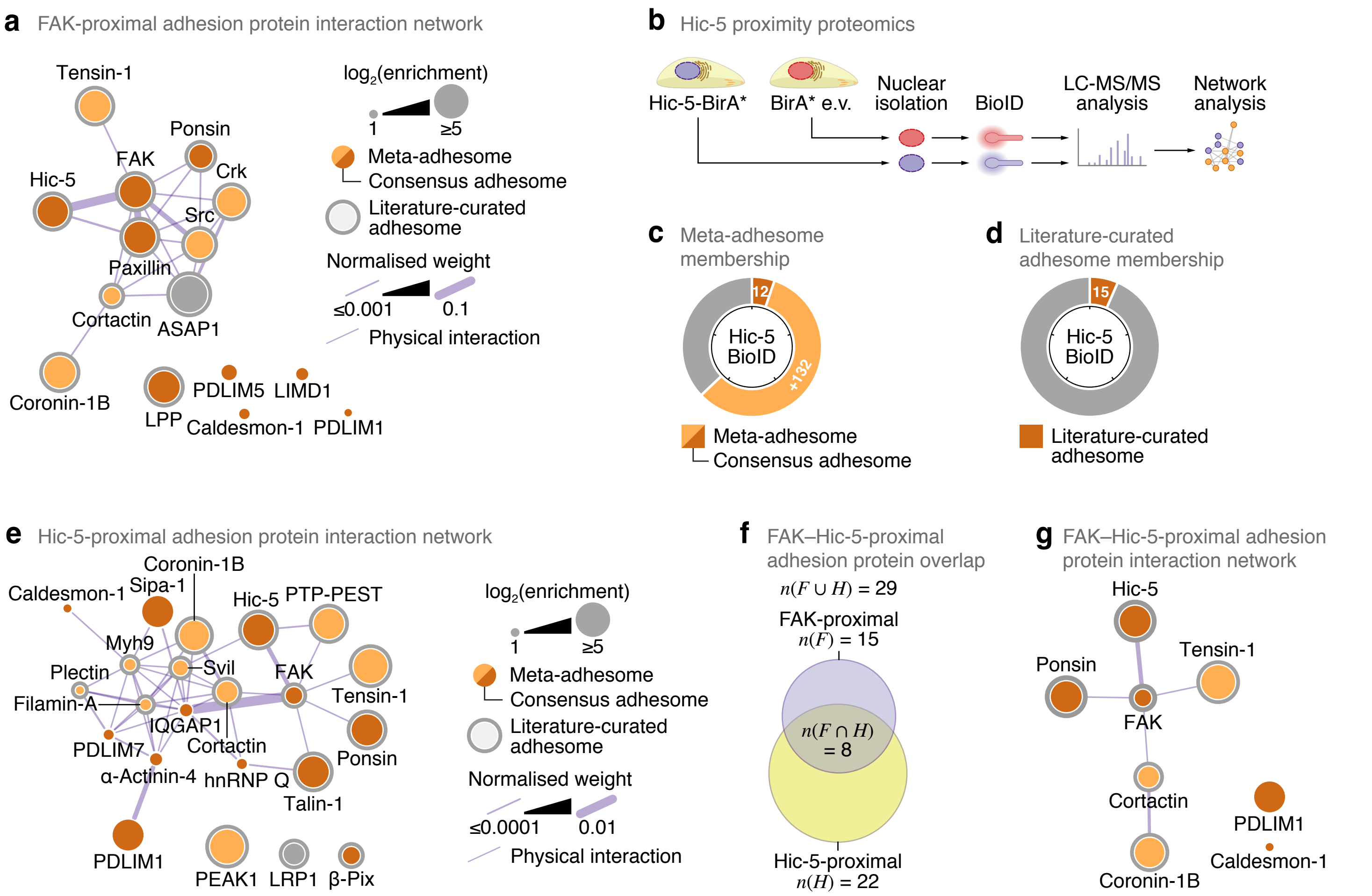


**Supplementary Figure 5.** Mass spectrometric analyses of the FAK- and Hic-5-proximal nuclear interactomes. (**a**) Interaction network analysis of specific FAK-proximal nuclear proteins present in the consensus or literature-curated adhesomes. (**b**) Workflow for mass spectrometric characterisation of Hic-5-proximal nuclear proteins in SCC cells. (**c**, **d**) Proportions of specific Hic-5-proximal proteins ($P < 0.05$, Student's $t$-test with FDR correction) quantified in the meta-adhesome, including the consensus adhesome (**c**), and the literature-curated adhesome (**d**). Tick marks indicate 20% increments. (**e**) Interaction network analysis of specific Hic-5-proximal nuclear proteins present in the consensus or literature-curated adhesomes. Myh9, myosin-9 (also known as myosin-IIa); Svil, supervillin. (**f**) Area-proportional Euler diagram of numbers of consensus or literature-curated adhesome proteins identified in FAK- and Hic-5-proximal nuclear interactomes. Cardinalities of sets are defined, where $F$ = FAK-proximal and $H$ = Hic-5-proximal nuclear adhesion proteins. (**g**) Interaction network analysis of the intersection of specific FAK- and Hic-5-proximal nuclear proteins present in the consensus or literature-curated adhesomes. Key as for **e**. For **a**, **e** and **g**, networks were constructed from reported physical protein interactions (edges) and weighted by linear regression according to evidence of protein co-association in the context of that network. Proteins are clustered according to connectivity; unconnected proteins also shown.
